# Evolution promotes transition-state-like conformations with enhanced basicity and electric fields in the substrate complex of a designer enzyme

**DOI:** 10.64898/2026.09.09.750330

**Authors:** Abbie Lear, H. Adrian Bunzel, Adrian J. Mulholland

## Abstract

Enzymes can be computationally designed for an increasing range of reactions, yet the catalytic efficiencies achieved are typically well below those of natural biocatalysts. Directed evolution can narrow this gap, often improving activity through mutations with non-obvious effects on catalysis. Here, we dissect how directed evolution improved a *de novo* designed Kemp eliminase by comparing the ground-state and transition-state ensembles of the designed and evolved variants. Extensive molecular dynamics simulations and QM/MM reaction barrier calculations identify subtle but important differences that modulate reactivity. Simulations show that reaction proceeds via a specific transition-state-like conformation, characterized by desolvation of the catalytic base and an organized active-site electric field. Barriers correlate strongly with the reaction energy of proton transfer, indicating that desolvating the catalytic base in the ligand-bound complex raises its proton affinity and thereby lowers the barrier. Evolution enriches the ground-state ensemble in reactive conformations, minimizing the reorganization required to reach the transition state. After evolution, formation of the transition-state-like conformation causes less disruption to the protein, which maintains coordinated motions. This is consistent with the emergence of a negative activation heat capacity with evolution. Efficient catalysis requires a ground-state ensemble that resembles the transition state, an objective that enzyme design should pursue alongside geometric and electrostatic complementarity for the transition state.

## INTRODUCTION

Enzymes are highly active and specific catalysts that work under mild conditions, making them attractive for sustainable industrial processes.^1–3^ Computational protein design, driven by deep learning, increasingly yields tailor-made enzymes to fill the gaps in the biocatalytic toolbox.^4–8^ Despite recent successes, designer enzymes often fall short of the catalytic power of their natural counterparts. Nonetheless, designed biocatalysts are typically highly amenable to directed evolution, and can be substantially improved through iterative rounds of mutagenesis and screening.^9–11^ Current computational design protocols therefore lack some important factors and the mutations selected during evolution may point to missing design features.^12^ Moreover, understanding the effects wrought by evolution can reveal fundamental insights into how enzymes work. Encoding these insights into design algorithms should improve enzyme design, accelerate biocatalyst engineering, and help access reactions and processes that are currently out of reach.

The theozyme approach is a central paradigm in enzyme design.^13,14^ Theozymes are minimal models typically comprising the chemical transition state (TS) and its surrounding catalytic residues. Structure-based enzyme design generally aims at embedding the theozyme geometry into a protein scaffold. Geometry, however, is only a proxy for catalytic activity. Catalysis originates from the electrostatic environment of the active site, which lowers the reaction barrier and, in doing so, often draws the substrate and catalytic residues into reactive arrangements.^15^ Enzymes gain much of their advantage over reactions in solution through preorganization, in which electrostatic interactions with the substrate are already arranged in the ground state (GS) to stabilize the chemical TS or reactive intermediates. Enzyme design algorithms often overlook preorganization and electrostatics.^14^ Even recent approaches that explicitly assess conformational ensembles typically assess preorganization by catalytic geometry rather than electrostatic features.^4,16^ Establishing which conformations within the GS ensemble are reactive and electrostatically preorganized, and what causes an enzyme to populate them, may reveal rules for designing preorganization.^12,17^

The Kemp elimination, the base-catalyzed deprotonation and ring-opening of benzisoxazoles, is a single-step benchmark reaction in enzyme design (Fig. 1a).^18–22^ The *de novo* enzyme studied here, 1A53-2, was designed by building an active site into a TIM barrel scaffold, giving moderate turnover numbers (*k*_cat_ = 0.0058 s^-1^).^23^ Subjecting 1A53-2 to directed evolution yielded 1A53-2.5, in which six mutations raise activity by more than three orders of magnitude (*k*_cat_ = 10 s^-1^, Fig. 1b+c).^11^ How do these mutations enhance activity, and what did design miss that evolution added? Kinetics offer a first indication, because evolution unexpectedly introduced an optimum in the temperature-rate profile, departing from the linear Arrhenius behavior of the designed variant.^11^ The curvature is not due to protein unfolding or instability, nor to a change in the rate-limiting step or mechanism. Previous MD simulations indicated that this curvature reflects the introduction of a negative activation heat capacity 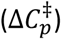 in 1A53-2.5, arising from the selective rigidification of the TS compared to the GS ensemble.^11,24^ Rigidification is associated with a network of correlated motions linking the active site and protein scaffold. While the apparent 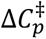 is a signature of improved catalysis, the origins of this behavior and the link to catalysis remain unclear, and this is actively debated.^25–28^

**Fig. 1.**
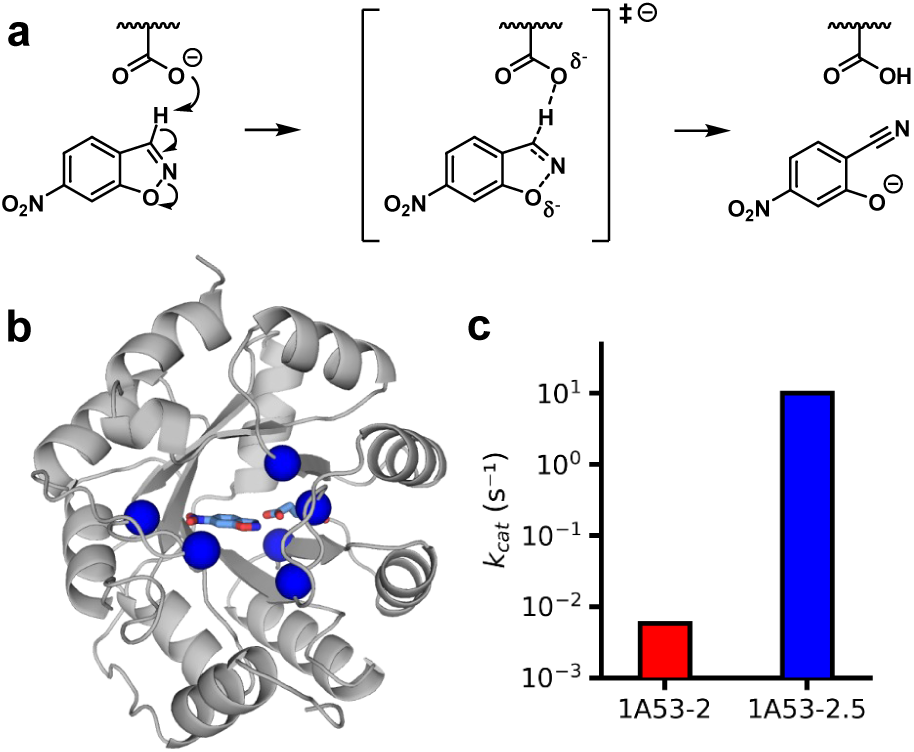
Evolution of a designer enzyme enhances enzyme activity. **a)** Base-catalyzed Kemp elimination**. b)** Structure of 1A53-2 with the six mutations yielding improved catalysis shown in blue. **c)** 1A53-2 was improved by more than 3 orders of magnitude through directed evolution, yielding 1A53-2.5.^11^

Here, we dissect the differences in reactivity between 1A53-2 and 1A53-2.5 using extensive MD simulations and QM/MM MD reaction profile calculations. By comparing substrate-bound ensembles with those generated with the chemical TS, we assess the presence of TS-like conformations in the GS ensemble. We find that evolution selectively enriched electrostatically preorganized, TS-like conformations with lower reaction barriers. Our work suggests that enzyme design should expand beyond theozyme geometry and additionally select for electrostatic preorganization, enriching the GS ensemble in reactive, preorganized conformations.

## RESULTS

### Evolution shifts the GS ensemble towards TS-like conformations

To assess how evolution changes the conformational ensemble, we performed multi-replicate MD simulations of 1A53-2 and 1A53-2.5 each bound to either the chemical GS or TS (15 × 500 ns = 7.5 μs per complex, Fig. S1). Our earlier simulations of this system required distance restraints between the ligand and the catalytic base.^25^ Here, we developed and tested new ligand parameters by deriving partial charges in the presence of the catalytic base to more accurately capture substrate polarization. The resulting ligand binding poses are substantially more stable, with distance restraints active in only 1.4% of frames for 1A53-2-GS (Fig. S1). Still, the active sites differ markedly between variants, with the GS and TS ensembles diverging in 1A53-2 but closely resembling one another in 1A53-2.5 (Fig. 2a,b). Average ensemble structures reveal that the ligand and catalytic base adopt a similar orientation in the GS and TS complexes of 1A53-2.5, whereas they are substantially reoriented in 1A53-2 (GS-TS RMSD 0.40 vs 1.22 Å, Fig. 2a). Principal component analysis of the ligand and base coordinates gives the same picture, with the 1A53-2 ensembles spanning a broad range along PC1 and overlapping by only 45 %, against 79 % for the narrower 1A53-2.5 distributions (Fig. 2b). Ligand and base distances and dihedrals show the same trend, with smaller differences between the GS and TS distributions for 1A53-2.5 (0.27 Å and 2.9 °) than 1A53-2 (0.54 Å and 39.2 °). (Fig. S2b-c). Together, these results indicate that evolution shifted the GS ensemble towards conformations sampled in the TS ensemble.

**Fig. 2.**
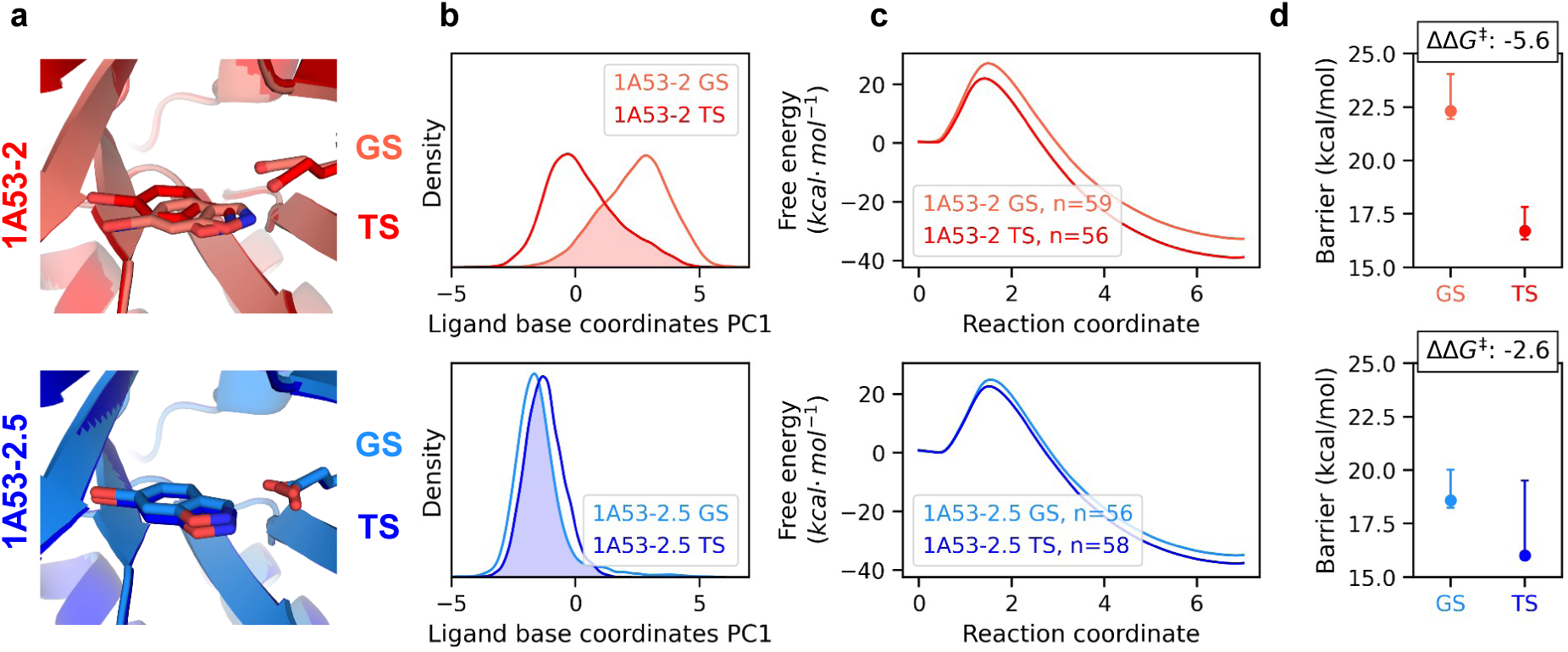
Directed evolution preorganises the enzyme-substrate complex, making it more TS-like. **a)** Average structures from MD simulation of the GS and TS ensembles are more similar in 1A53-2.5 than in 1A53-2. **b)** Principal component analysis of ligand and catalytic base coordinates shows increased GS–TS overlap after evolution (1A53-2: 45% overlap; 1A53-2.5: 79% overlap; PC1 covers 39% variance). **c)** Mean free energy profiles of the GS and TS ensembles of 1A53-2.5 are similar (blue), whereas they are different for 1A53-2 (red, PM6/CHARMM36). **d)** Barriers calculated for GS (darker color) and TS (lighter color) ensembles show that the GS ensemble becomes more reactive and TS-like after evolution (error bars indicate bootstrapped uncertainties of the Boltzmann average from 60 barriers per ensemble).

To determine how structural preorganization affects the chemical reaction, ≈60 quantum mechanics/molecular mechanics (QM/MM) reaction free-energy profiles were calculated per ensemble. Free-energy profiles were calculated using the adaptive string method, which iteratively refines the reaction pathway across multiple collective variables and projects it onto a single reaction coordinate (Tab. S1).^29^ The resulting barrier difference between the GS ensembles of 1A53-2 and 1A53-2.5 is close to the experimental activity difference (ΔΔ*G*^‡^ calculated: 3.7 kcal mol^-1^, experiment: 4.4 kcal mol^-1^, Fig. 2c+d). Absolute barriers are overestimated by ≈2 kcal mol^-1^ relative to experiment, reflecting limitations of PM6 which was chosen after preliminary testing of semi-empirical methods to balance accuracy with computational efficiency. In both variants, structures from the TS ensemble exhibit lower barriers than those from the GS ensemble. The gap between them, however, is far smaller for 1A53-2.5 than for 1A53-2 (2.6 kcal mol^-1^ vs 5.6 kcal mol^-1^). The TS ensembles of both variants therefore react with similar barriers (Tab. S2). Evolution apparently did not only increase the reactivity of the most reactive states, but enriched the GS ensemble of 1A53-2.5 in these catalytically competent, TS-like conformations.

### Electrostatic Preorganization Enhances Activity

Reaction barriers are often correlated with reaction free energy in a linear free-energy relationship, indicating effects that act along the entire reaction coordinate rather than specifically on the chemical TS. We therefore compared Δ*G*^‡^ with the reaction energy (Δ*G*, the energy difference between the GS and product state, Fig. 3a). The reaction energy correlates with the reaction barrier across all reaction profiles independent of ensemble (R^2^ = 0.78, Fig. 3b). The slope is close to unity, so that a change in Δ*G* is passed on almost entirely to the barrier. Differences in Δ*G* are therefore sufficient to account for their differences in reactivity. Consistent with this, the TS position shifts earlier along the reaction coordinate as the reaction becomes more exergonic (R^2^ = 0.22, Fig. S3c), as expected from the Hammond postulate. Comparing ensemble averages, the reaction energy is more exergonic in the TS ensembles (1A53-2: −38.7 ± 0.5 kcal mol^-1^, 1A53-2.5: −37.5 ± 0.5 kcal mol^-1^) than in the GS ensembles (1A53-2: −32.8 ± 0.6 kcal mol^-1^, 1A53-2.5: −35.0 ± 0.6 kcal mol^-1^). As observed for Δ*G*^‡^, the Δ*G* difference between the GS and TS ensembles becomes smaller after evolution (5.5 vs. 2.5 kcal mol^-1^). The smaller GS–TS gap in 1A53-2.5 reflects a GS ensemble already enriched in reactive TS-like conformations.

**Fig. 3.**
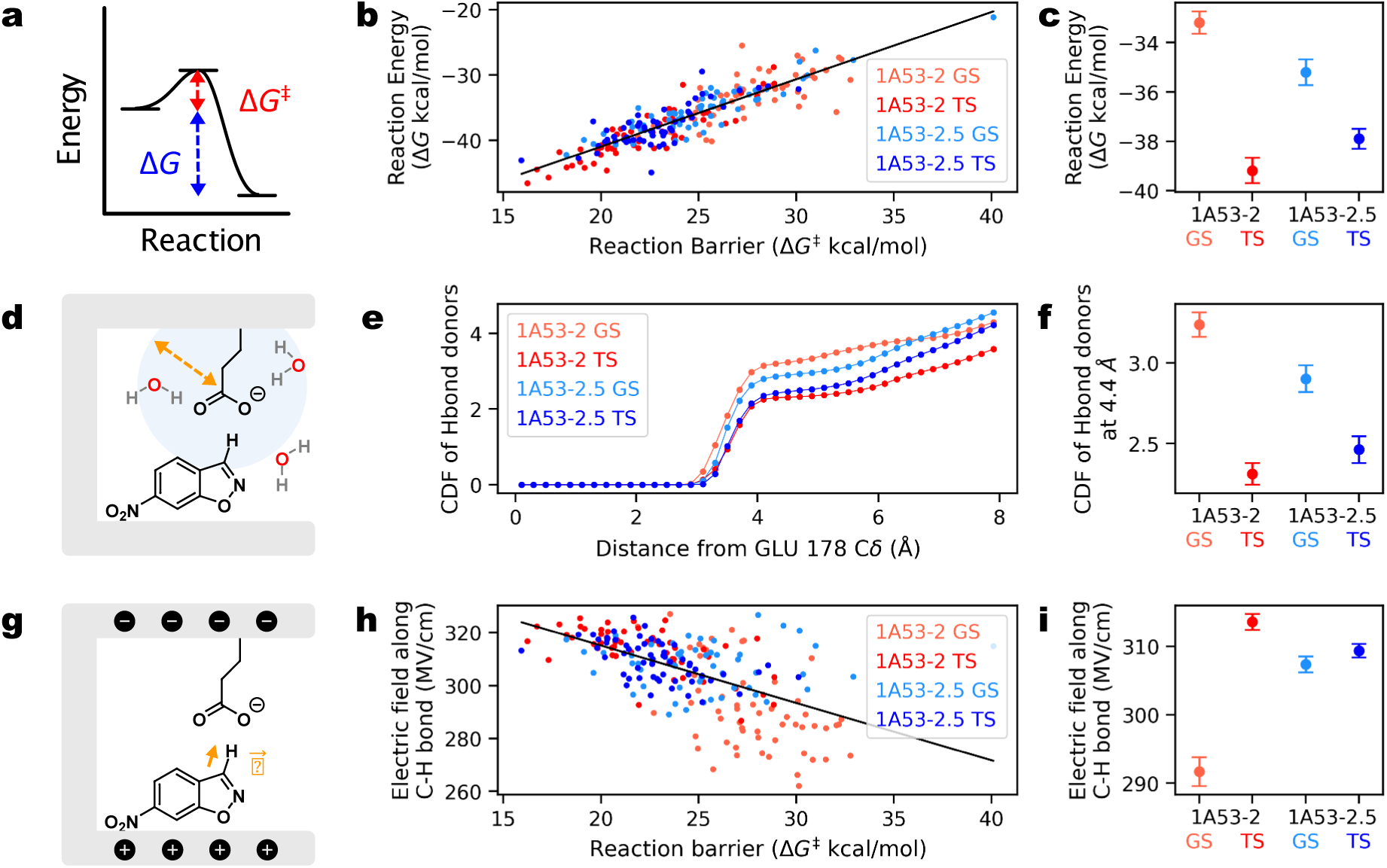
Evolution boosts reactivity by reducing hydrogen bonding to the catalytic base. **a)** Comparison between reaction barrier (Δ*G*^‡^) and reaction energy (Δ*G*). **b)** Δ*G* plotted against Δ*G*^‡^ for all PM6/CHARMM36 reaction simulations for the GS and TS ensembles of 1A53-2 and 1A53-2.5. Linear fit gives R^2^ = 0.78 and p = 2 × 10^-77^. **c)** Average reaction energy over the four ensembles. **d)** Cumulative distribution functions were calculated for hydrogen bond donors to the carboxylate carbon of the catalytic base (Cδ of Glu178). **e)** Average cumulative radial distribution functions were calculated from equilibrium QM/MM MD simulations **f)** Average sum of hydrogen bond donors up to 4.4 Å for each ensemble. **g)** Electric fields were calculated along the scissile C-H bond. **h)** Electric fields plotted against reaction barrier for all reaction simulations. Data are calculated from structures ± 0.5 amu^1/2^ Å from the transition state of the reaction free energy profiles shown in Fig. 2. Linear fit gives R^2^ = 0.34 and p = 3 × 10^-22^. **i)** Electric fields averaged for each ensemble (error bars in c,f & i show the standard error of the mean over all reaction profiles for each ensemble).

Reactivity in the Kemp eliminases is apparently set by Δ*G*, raising the question of how the protein environment changes the reaction energy. A crucial factor in determining Δ*G* will be the reactivity of the catalytic base so we examined its hydrogen-bonding environment. Across the QM/MM equilibrium trajectories, conformations with fewer hydrogen-bond donors show lower barriers (<20 kcal mol^-1^: 2.09 ± 0.11 H-bonds; >20 kcal mol^-1^: 2.80 ± 0.05; Fig. S3a). Comparing variants, the GS ensemble of 1A53-2.5 contains fewer hydrogen-bond donors Fig. 3e), and the difference in solvation between the GS and TS ensembles becomes much smaller after evolution (1A53-2: 0.93, 1A53-2.5: Fig. 3c,f).

The structural and solvation changes described above will also have electrostatic effects, as desolvation and repositioning of the base are expected to strengthen the electric field acting on the scissile bond.^17^ Using FieldTools, we calculated the electric field projected by the protein onto the scissile C–H bond for structures close to the TS from each QM/MM reaction simulation (Fig. 3g).^30^ The field along the C–H bond has some correlation with both Δ*G*^‡^ (R^2^ = 0.34, p = 3 × 10^-22^, Fig. 3h) and Δ*G* (R^2^ =0.32, p=1.6 × 10^-20^, Fig. S3b). Conformations with fewer hydrogen-bond donors to the carboxylate exhibit stronger fields (Fig. S3d), which suggests that the electric field affects the reactivity of the base. Evolution increases the average field of the GS ensemble from 292 ± 2 MV cm^−1^ to 307 ± 1 MV cm^−1^, close to values in the TS ensembles themselves (314 ± 1 MV cm^−1^ and 309 ± 1 MV cm^−1^, Fig. 3i). This shift suggests that evolution lowers the reaction barrier by preorganizing the GS ensemble towards conformations with stronger catalytic electric fields.

### Conformational dynamics and the evolution of non-Arrhenius behavior

Evolution of 1A53-2 unexpectedly altered its activity-temperature dependence and introduced curved non-Arrhenius behavior.^11^ This behavior has been explained by macromolecular rate theory,^24^ indicating a negative activation heat capacity 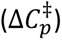 in the evolved variant.^11^ This implies reduced energetic fluctuations of the TS relative to the GS. Previous simulations of 1A53-2.5 showed this and an increase in correlated dynamics in the TS versus the GS, not observed in 1A53-2.^25^ These simulations employed a distance restraint between the ligand and base, which could affect the calculated 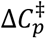.^24,26–28,31^ The updated ligand parameters developed here alleviate this restraint (Fig. S1). Here, we also include solvent effects using an implicit solvent model. These calculations give a negative 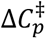 for 1A53-2.5 (−0.09 ± 0.08 kJ mol^-1^ K^-1^) and a positive 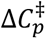 (0.15 ± 0.08 kJ mol^-1^ K^-1^) for 1A53-2 of similar orders of magnitude to the experiment (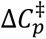 = −0.28 kJ mol^-1^ K^-1^ for 1A53-2.5) (Fig. 4a, Fig. S4a). The difference between the activation heat capacities of the two variants is comparable to the experimental difference (calculated: 0.23 kJ mol^-1^ K^-1^, experimental: 0.28 kJ mol^-1^ K^-1^).

**Fig. 4.**
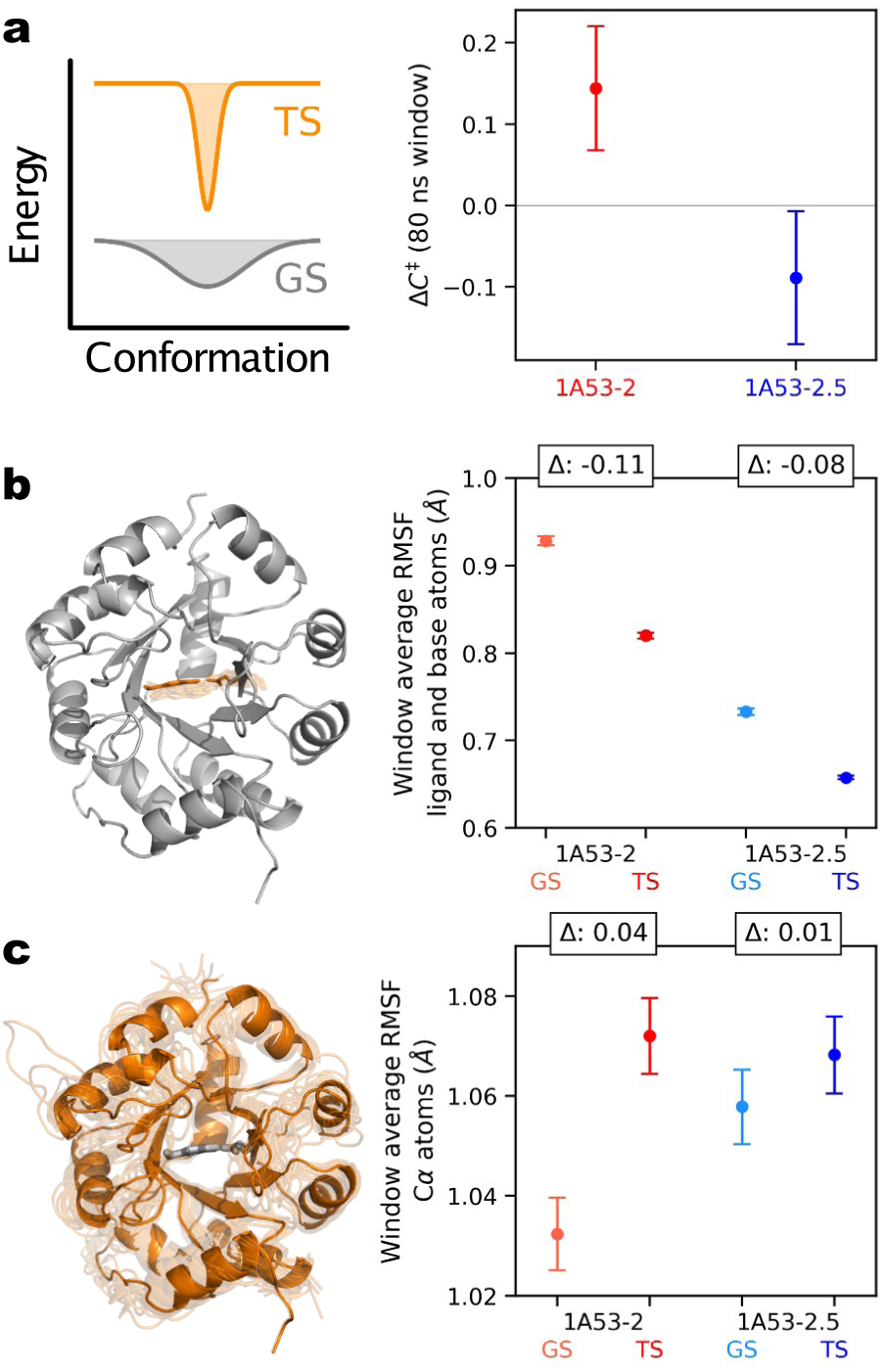
Local and global changes in conformational dynamics. **a)** The activation heat capacity was calculated using Eq. 1 with an 80 ns window average approach^25,35^ from the difference in energetic fluctuations between the GS and TS ensembles. Energies are calculated using a GBSA implicit solvent after removal of the explicit solvent atoms. Error bars represent bootstrap confidence intervals from replicate simulations with replacement. **b-c)** Average root mean squared fluctuation (RMSF) of the ligand and base atoms **(b)** and all protein Cα atoms **(c)** using an 80 ns window for the GS and TS ensembles of both enzyme variants. Error bars represent the standard error of the mean.

The apparent negative 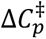 in 1A53-2.5 reflects a difference in energetic fluctuations between the TS and GS ensembles, but does not reveal which motions are involved. To obtain a structural picture, we calculated root-mean-square fluctuations (RMSFs) from MM MD simulations. Both enzymes show a decrease in RMSF values between the GS and TS ensembles for the ligand and base atoms (−0.11 Å vs −0.08 Å for 1A53-2 and 1A53-2.5, Fig. 4b), which reflects the organization of the active site associated with reaction. However, in the case of 1A53-2, active site tightening is accompanied by an increase in backbone flexibility (Cα RMSF 0.04 Å vs 0.01 Å, Fig. 4c, Fig. S4b). This increase in flexibility indicates that organizing the active site to populate reactive TS-like conformations destabilizes 1A53-2, which is likely associated with an energetic penalty and increased energetic fluctuations. In contrast, the backbone RMSF remains essentially unchanged in 1A53-2.5 between the GS and TS ensembles. Evolution reduces the difference in backbone structural fluctuations between the GS and TS ensembles which is consistent with reduced energetic fluctuations in the TS complex that give rise to the apparent negative 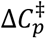. In the evolved variant, the reaction can occur without significant reorganization or loss of interactions in the protein because the protein is better organized for reaction (including carboxylate desolvation). This provides a rationale for why the increase in correlated dynamics apparent after evolution is connected to better catalytic power.

Taken together, our QM/MM and fluctuation results point to a common origin of the differences in reactivity and dynamics between the two variants. The QM/MM results show that reactive TS-like conformations are enriched after evolution. Efficient reaction requires the GS ensemble to reach one of these conformations, narrowing the accessible conformational space. This narrowing probably causes the negative 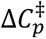 in 1A53-2.5. By contrast, 1A53-2 is less preorganized and must rearrange to reach reactive TS-like conformations. This reorganization destabilizes the surrounding protein, reflected in increased RMSF values, which likely prevents the conformational narrowing and reduction in energetic fluctuations necessary for a negative 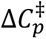. While we recognize that the fluctuation method has important limitations and caveats for calculating heat capacity differences from simulations, we nonetheless observe significant differences between the two systems and systematic errors are cancelled by subtraction.^32^ We also note that our observations of reduced fluctuations between the GS and TS ensembles could alternatively be described in terms of a two-state model.^31,33,34^ A shift from a broad, flexible ensemble to a narrower, more rigid one is qualitatively similar to a population shift between an inactive and an active state. We thus suggest that, where non-Arrhenius kinetics arise from such a population shift rather than from a rate-limiting conformational inactivation, binary and 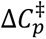-based models may often capture the same underlying effects that cause non-Arrhenius kinetics.

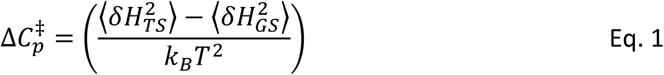

## DISCUSSION

To understand how directed evolution improves enzyme catalysis, we compared a computationally designed Kemp eliminase with an evolved variant using extensive MD simulations and QM/MM reaction free-energy calculations. Simulations of the GS ensemble reproduce the activity gain achieved by evolution, and comparison with the TS ensembles reveals that evolution enriched the GS ensemble with low-barrier, TS-like conformations (Fig. 5). Both variants react with similar barriers once the TS ensemble is reached, so evolution predominantly increased the population of the reactive state. Reactive conformations exhibit precise structural organization and reduced reaction barriers in part due to active site desolvation, leading to catalytically superior electric fields and a more reactive base. In the designer enzyme, reaching these conformations requires substantial reorganization that disrupts the protein and increases its energetic fluctuation. In contrast, the evolved variant accesses the same conformations without such a penalty, which likely enables its apparent negative 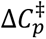.

**Fig. 5.**
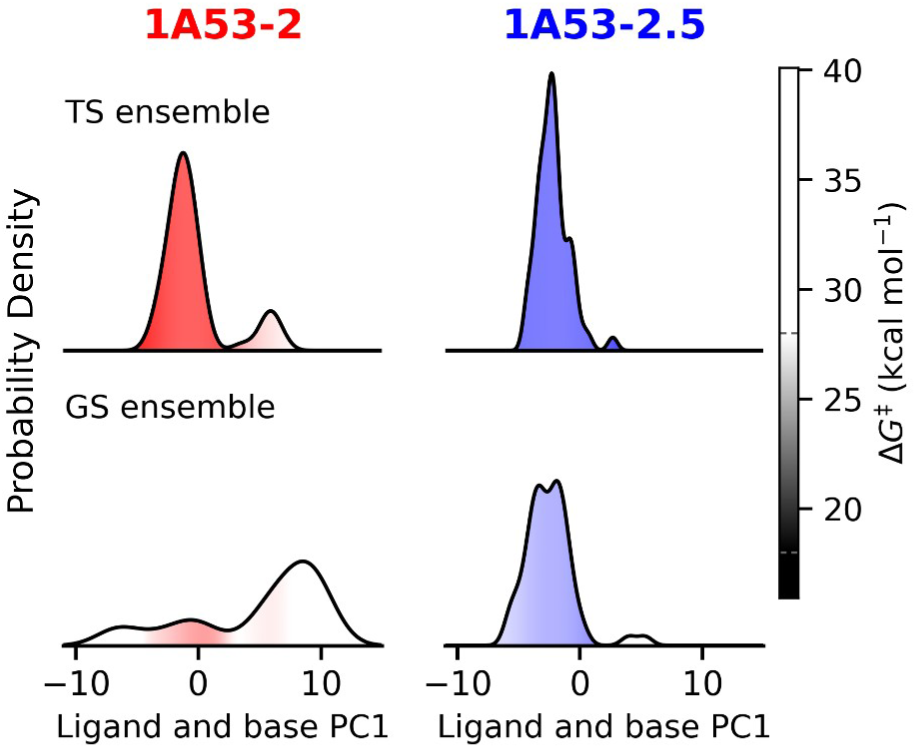
Evolution enriched reactive TS-like states in the GS ensemble. GS and TS ensembles of 1A53-2 (red) and 1A53-2.5 (blue) shown as kernel densities estimates along PC1 (45% variance) of the ligand and base coordinates. PCA was performed using ligand and base coordinates from frames ± 0.05 amu^1/2^ Å from the TS observed in QM/MM reaction barrier calculations and densities are colored by the calculated barrier using a local average (saturated, low barrier; white, high barrier).

The Kemp elimination is base-catalyzed, so the increased reactivity of the base in the GS ensemble of 1A53-2.5 implies that the proton affinity of the base has increased. However, our previous experiments show that evolution lowers the base p*K*_a_ from 8.7 to 5.4, an apparent contradiction that is resolved by the different environments of the apo and substrate-bound states.^11^ The experimental p*K*_a_ reports on the apo state, where the higher value of 8.7 in 1A53-2 would in principle give the more reactive base but leaves it largely protonated at pH 7. Lowering the p*K*_a_ keeps the base deprotonated, as required for base catalysis, at the cost of reactivity. Substrate binding then displaces the active-site water, increasing proton affinity. Evolution appears to have optimized both states, lowering the apo p*K*_a_ while arriving at a more desolvated, and therefore more reactive, base in the substrate complex.

Desolvation of a carboxylate base and electric field catalysis are recurring catalytic strategies in natural enzymes.^36,37^ In ketosteroid isomerase (KSI)^38^ and triosephosphate isomerase (TIM)^39^, simulations show that desolvation of the base by active-site closure is required for efficient reaction. In aromatic amine dehydrogenase, reactivity is likewise sensitive to the hydrogen-bonding environment of the base.^40^ Comparison of OXA-48 and OXA-163 class D β-lactamases shows that the latter’s improved activity against cephalosporins is due to a small reduction in hydration of the active-site base.^41^ Evolution of 1A53-2 therefore converged on a mechanism that natural enzymes widely exploit. Removing water does not by itself create a better environment. Enzymes replace water with preorganized polar groups that stabilize the developing charge more effectively,^42^ which is precisely what happened during evolution of 1A53-2. Arranging polar groups around a charge that has yet to develop leaves the catalytic residues in locally destabilized, energetically frustrated environments.^43,44^ Energetic frustration and reactivity are inseparable, because a site relaxed around the substrate would have to reorganize to reach the transition state, a cost preorganization pays in advance. Most design algorithms, however, aim to maximize stability. Creating a reactive active site poses the significant challenge of introducing the local destabilization these algorithms are written to eliminate.

Electrostatic preorganization of 1A53-2.5 emerged through evolution of the conformational ensemble. Evolution is known to act on designer enzyme ensembles in various ways,^45,46^ frequently stabilizing alternative substrate conformations,^20,47^ or changing the catalytic residue.^48–50^ Evolutionary fine-tuning in 1A53-2 is more subtle than restructuring the whole active site and instead organizes conformations within a similar binding mode towards more reactive, TS-like state. TS ensemble organization has been demonstrated in other designer enzymes, for instance, through crystallography of transition-state analog complexes of Kemp eliminase HG3.17.^16,51^ Likewise, our earlier study of 1A53-2.5 described organization of the TS ensemble arising through conformational narrowing and the emergence of a dynamical network.^25^ Here, we complement these studies by demonstrating that evolution enriches the GS ensemble in reactive conformations.

Current design pipelines based on generative deep learning models can now create *de novo* catalysts with moderate success.^4–6,8,52,53^ While state-of-the-art pipelines typically focus on stabilizing the chemical TS, enzyme design should also directly target the GS ensemble and score its similarity to the TS ensemble. Established approaches typically optimize the fit of a single structure to the chemical TS, and where electrostatics are targeted, catalytic fields are evaluated on static or ensemble-averaged structures.^17,54,55^ Ensemble-aware approaches have begun to address conformational fluctuations through crystallographic ensembles,^16,56^ multi-state design,^57^ and ML-generated conformational ensembles.^58^ However, ensembles are typically assessed for the apo enzyme or the TS complex. Moreover, preorganization is typically judged by the flexibility of catalytic groups rather than by the reactivity of the sampled conformations. While generating GS- and TS-bound ensembles requires only suitable ligand parameters, accurately and quickly building ensembles remains challenging, as does capturing active-site hydration and performing multi-state optimization. It is also essential to note that conformations that allow for efficient substrate binding and product release are unlikely to be those in which optimal catalytic organization is achieved, but both conformations of the substrate complex must be accessible for overall rapid turnover.^59^ Frameworks integrating design algorithms, ensemble generators, and activity predictors in a multi-objective optimization loop, such as AI.zymes,^54^ will likely be needed to efficiently implement these complex design objectives.

## CONCLUSION

Evolution optimizes 1A53-2 by shifting the GS ensemble towards reactive, TS-like conformations. This shift is accompanied by both a stronger catalytic electric field and a less solvated catalytic base. While current design efforts largely focus on geometric complementarity to the chemical TS, our work suggests that targeting the electrostatic and conformational preorganization of the GS ensemble alongside TS complementarity will yield superior biocatalysts.

## METHODS

### Molecular Dynamics Simulations

MD simulations were run using AMBER22.^60^ Octahedral solvent boxes with 10 Å between the protein and box edge were set up using CHARMM-GUI^61,62^ Ligand charges were generated by the RED Server,^63^ which uses RESP^64^ fitting using DFT (B3LYP,^65^ 6-311+G(d,p), SCRF=water, Gaussian16^66^) derived geometries (See Fig. S1). Input structures from crystal structures 3NZ1^23^ (1A53-2.0) and 6NW4^11^ (1A53-2.5) were minimized and heated with backbone restraints (all Cα atoms and ligand and base atoms with a force constant of 100 kcal mol^-1^ Å^-2^). Restraints were released over 5 stages of 0.1 ns with decreasing force constants (20,8,4,2,1 kcal mol^-1^ Å^-2^), followed by free equilibration in the NVT ensemble for 2 ns based on previous protocols.^25^ Production was run in the NPT ensemble using the Berendsen barostat with a coupling constant of 1 ps and Langevin dynamics for temperature control with a collision frequency of 1 ps^-1^ at 298 K for 500 ns. Two harmonic restraints were used throughout to ensure the ligand remained in the active site: C of the scissile bond in the ligand to Cα of the catalytic base was restrained to 8.5 Å, and the sidechain dihedral angle of Trp210 was restrained between 120° and 180° with a force constant of10 kcal mol^-1^ rad^-2^ for both restraints. See Fig. S1 for further details about the updated restraints and parameters.

### Adaptive string method

The adaptive string method^29^ (ASM) is an enhanced sampling technique that allows efficient, accurate simulation of enzyme catalysed reactions through a flexible definition of the reaction coordinate. Unlike multidimensional umbrella sampling, it does not scale with the number of collective variables as it only samples along a one-dimensional path. Sampling around the initial guess for the reaction pathway allows the true minimum energy pathway to be found. Here, we used 32 nodes and four collective variables detailed below (Tab. S1). Four structures from each 500 ns MD run (timepoints 150, 250, 350, and 500 ns) were equilibrated with QM/MM (PM6/CHARMM36) simulations using AMBER22^60^ for 100 ps. The QM region included the ligand atoms and CH_2_CO_2_ of the catalytic base. The minimum free-energy path was optimized for 20 ps and then sampled for 100 ps. Structural analysis was performed using pytraj,^67,68^ sklearn^69^ and pymol.^70^

### Electric field calculations

Electric fields were calculated using FieldTools.^30^ FieldTools uses the point charges from the CHARMM36 force field to calculate the electrostatic potential of the surrounding protein at a particular point or along a vector using Coulomb’s law. The fields described here are calculated along the vector of the scissile C-H bond of the ligand. The ligand atoms are excluded from the calculation, so the calculated field effect only comes from the protein. The electric field values shown here (Fig. 3) are calculated from structures ± 0.5 amu^1/2^ Å from the highest energy point of the reaction profile calculated using QM/MM adaptive string calculations.

### Activation heat capacity calculations

Activation heat capacities are calculated using the difference in energetic fluctuation between the GS and TS ensembles (Eq. 1).^25,26,35,71^ To avoid the calculations being dominated by the contribution from the solvent, water is removed from MM MD structures. The energy of each frame is recalculated using the CHARMM36 force field. Solvation effects are approximated^72^ using the generalized Born solvent accessible method available in AMBER22 (igb=8).^73^ The energy is recalculated for all saved snapshots (50,000 per MD run) and the window average variance is calculated. See below for discussion on window size selection (Fig. S4). This variance is averaged across all simulations, and the difference in variance is calculated to determine the activation heat capacity.

## SUPPORTING INFORMATION

Additional data are described in Supplementary Tables S1 and S2 and Supplementary Figures S1-4.

## CORRESPONDING AUTHORS

H. Adrian Bunzel, Adrian J. Mulholland

## AUTHOR CONTRIBUTIONS

Conceptualization: AL, HAB, AJM; investigation, formal analysis, visualization: AL; writing – original draft: AL; writing – review and editing: AL, HAB, AJM; supervision: HAB, AJM; funding acquisition: HAB, AJM. All authors have given approval to the final version of the manuscript.

## ACKNOWLEDGEMENTS

This work is part of a project that has received funding from the European Research Council under the European Horizon 2020 research and innovation programme (PREDACTED Advanced Grant Agreement no. 101021207) to AJM. HAB thanks the SNSF (PZ00P3_208691), the Max Planck Society, and Max Planck Foundation for support. This work was conducted using the computational facilities of the Advanced Computing Research Centre, University of Bristol (http://www.bris.ac.uk/acrc/).

