## Supplementary Information for "Evolution promotes transition-state-like conformations with enhanced basicity and electric fields in the substrate complex of a designer enzyme"

**Contents**

1. Supplementary Tables 2

Tab. S1 | Collective variables used for reaction barrier calculations. 2

Tab. S2 | Thermodynamic data 2

2. Supplementary Figures 3

Fig. S1 | Ligand parameters and restraints do not significantly alter dynamics. 3

Fig. S2 | Structural features from MM MD simulations 4

Fig. S3 | Effects of hydrogen bonding on reaction barrier and electric field. 5

Fig. S4 | Further details of structural and energetic fluctuations. 6

### 1. Supplementary Tables

**Tab. S1 | Collective variables used for reaction barrier calculations.**

| Collective variable | Start value [a] | End value [a] |
|---|---|---|
| C – H distance | 1.07 Å | 3.20 Å |
| N – O distance | 1.43 Å | 3.60 Å |
| C – C – N angle | 110° | 180° |
| H – Cδ distance | 2.80 Å | 2.00 Å |

a) Values were selected based on DFT (B3LYP,[65] 6-311+g(d,p), SCRF=water, Gaussian16[66]) optimized geometries for the reactant and product states.

**Tab. S2 | Thermodynamic data**

| | 1A53-2 | 1A53-2.5 |
|---|---|---|
| $k_{cat}$ ($s^{-1}$)[a] | 0.0058 ± 0.0008 | 10 ± 1 |
| $\Delta G^{\ddagger}_{exp}$ (kcal $mol^{-1}$)[b] | 20.5 ± 0.1 | 16.1 ± 0.1 |
| $\Delta G^{\ddagger}_{calc}$ (kcal $mol^{-1}$)[c] | 22.3 (95% CI [21.9, 24.1]) | 18.6 (95% CI [18.2, 19.9]) |
| $\Delta G$ (kcal $mol^{-1}$)[d] | -33.2 ± 0.4 | -35.2 ± 0.5 |
| $pK_a$[a] | 8.7 ± 0.3 | 5.4 ± 0.1 |
| $\Delta G_{deprot}$ (kcal $mol^{-1}$)[e] | 2.3 ± 0.4 | -2.2 ± 0.1 |
| $K_m$ (µM)[a] | 1200 ± 300 | 710 ± 130 |

a) From reference [11].
b) Calculated from $k_{cat}$.
c) Calculated as Boltzmann average across reaction profiles from the GS ensemble generated in this work. Errors represent bootstrapped 95% confidence intervals.
d) Calculated as the average across reaction profiles from the GS ensembles generated in this work. Errors represent the standard errors of the mean.
e) Calculated from pKa at pH 7.

#### 2. Supplementary Figures

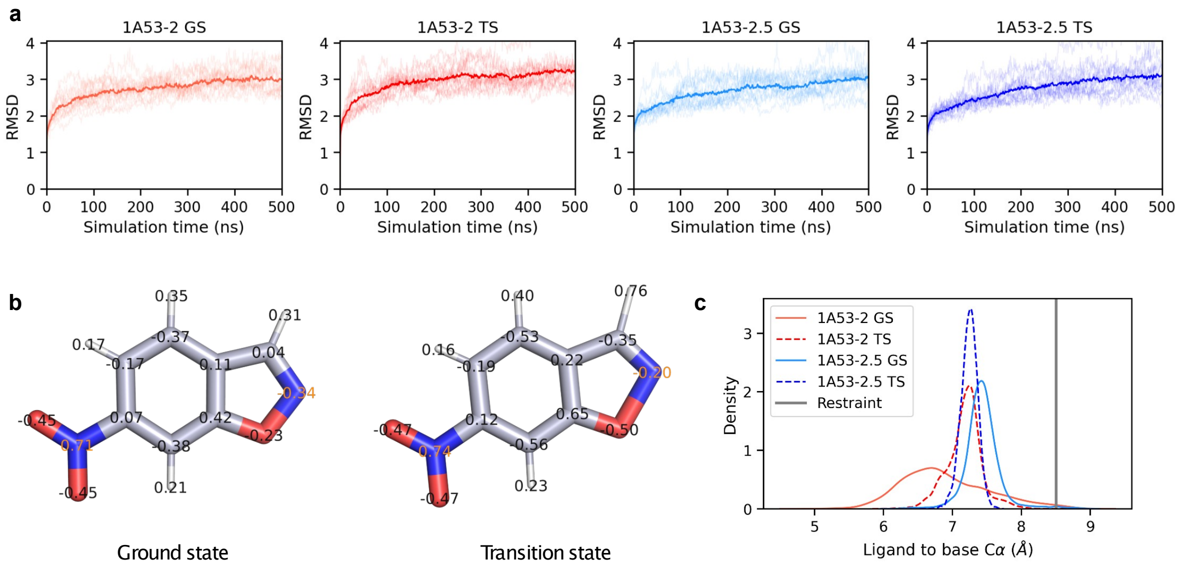


**Fig. S1 | Ligand parameters and restraints do not significantly alter dynamics.**

**a)** RMSDs for all 15 replicate MM MD simulations, for each of the four systems studied, with the average shown in bold. All MD analysis was performed on every 100$^{th}$ frame of the MD trajectories unless otherwise indicated. **b)** Ligand charges were derived using RESP[64] fitting from DFT (B3LYP,[65] 6-311+g(d,p), SCRF=water, Gaussian16[66]) derived geometries. Atomic charges are shown on the DFT-optimized structures. Nitrogen charges are shown in orange for clarity. Only the charge distribution was altered between the GS and TS simulations, as the geometric change is small. **c)** A distance restraint from the ligand to the catalytic residue was added, but only rarely activated for structures that show distances greater than 8.5 Å (1A53-2 GS: 1.4%; 1A53-2.5 G: 0.9%; 1A53-2 TS 0.04%; 1A53-2.5 TS: 0.0%).

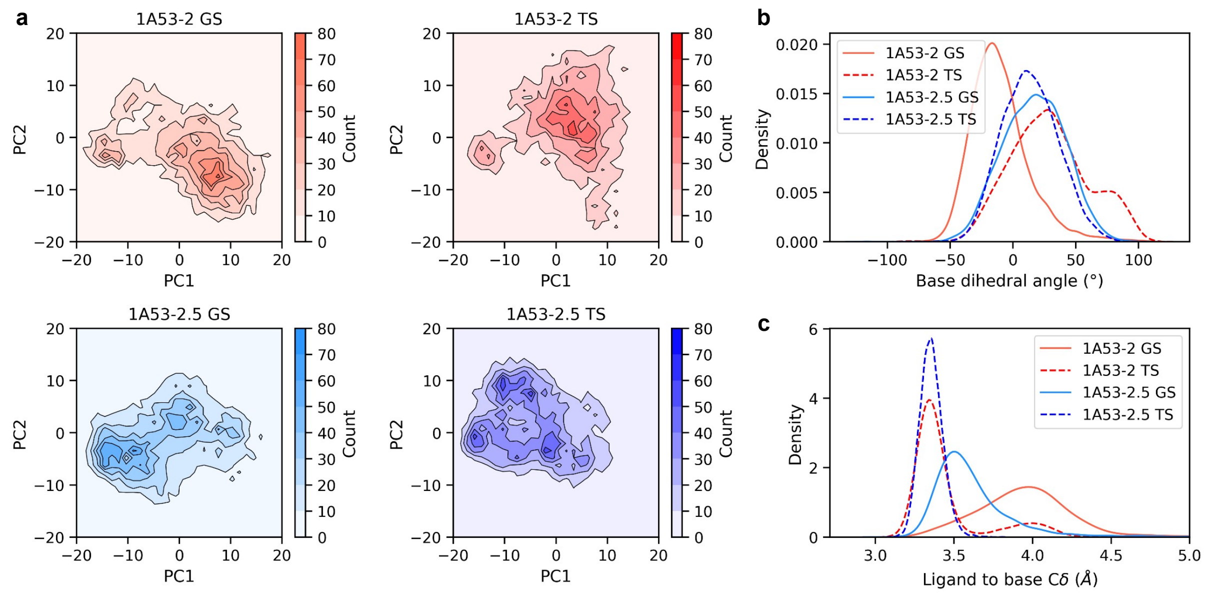


**Fig. S2 | Structural features from MM MD simulations**

**a)** The first and second principal components from PCA analysis of the protein backbone coordinates across all ensembles are plotted for each variant. Note the large difference between GS and TS for the designed variant, but a small difference for the evolved variant, indicating better overlap of the GS and TS ensembles after evolution. **b-c)** The distributions of the catalytic base dihedral (**b**) and the distance between the reacting carbon atom of the ligand and base Cδ (**c**) over the length of the simulations. Note the similarities between 1A53-2.5 GS and TS, but the difference between 1A53-2 GS and TS. The dihedral angle is calculated using the Cα, Cβ, Cγ, and Cδ atoms of the glutamate base.

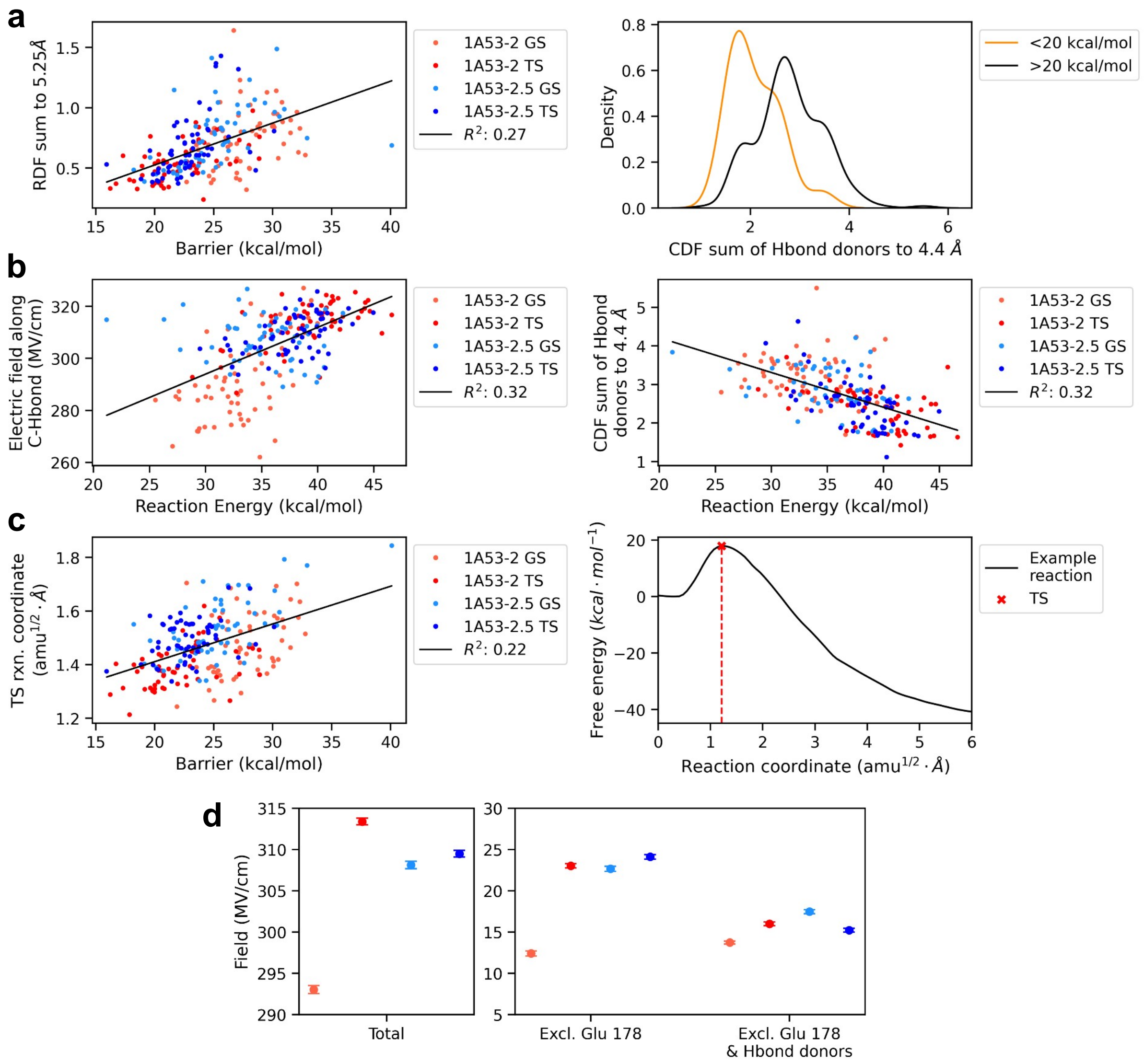


**Fig. S3 | Effects of hydrogen bonding on reaction barrier and electric field.**

**a)** Hydrogen bonding to the catalytic base has a significant effect on the barrier to reaction, as shown by the correlation between the reaction barrier and the sum of hydrogen bond donors within 5.25 Å of the catalytic base ($H_2O$ and Tyr157 for 1A53-2.5). Accordingly, ensembles with a barrier lower than 20 kcal/mol (orange) exhibit a lower count of hydrogen-bond donors (2.09 ± 0.11 H-bonds) within 4.4 Å than structures with a barrier greater than 20 kcal/mol (black, 2.80 ± 0.05 H-bonds). **b)** Both hydrogen bonding at the catalytic base and the electric field along the scissile bond can also be linked to reaction energy, as demonstrated by the correlation plots suggesting that all these effects arise together. **c)** The position of the TS on the reaction coordinate also correlates with the barrier to reaction. **d)** The electric field along the scissile bond is affected by hydrogen bonding: the largest differences in field between the four ensembles studied come from the catalytic base and residues that donate hydrogen bonds to the base ($H_2O$ for both variants and Tyr157 for 1A53-2.5).

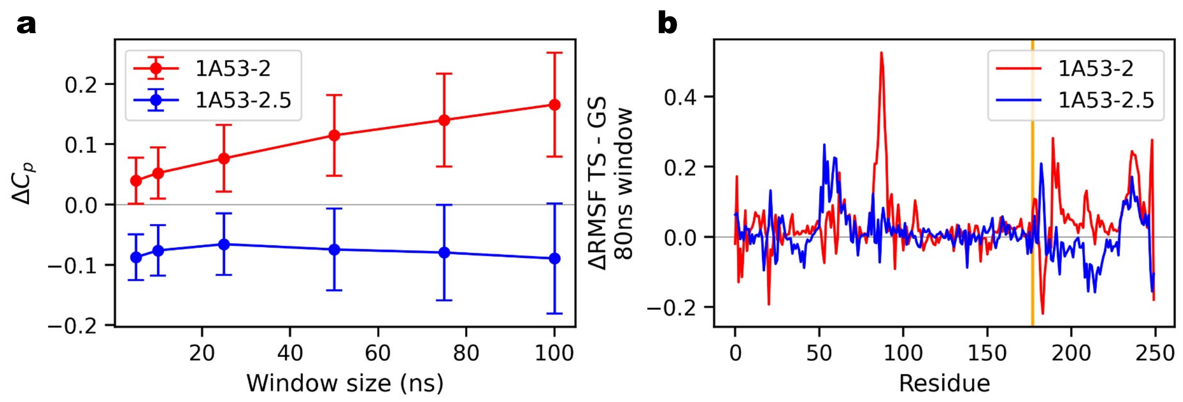


**Fig. S4 | Further details of structural and energetic fluctuations.**

**a)** Investigation of the correct size of window to use when calculating activation heat capacity. To remove the effects of large-scale conformational motion in the fluctuation calculations, the average fluctuation is computed using a moving-window approach. As in our previous work, an 80 ns window was used for further fluctuation analysis.[25,35] **b)** Average difference between GS and TS in RMSF per residue for 1A53-2.0 (red) and 1A53-2.5 (blue), with the position of the catalytic base shown in orange. Note the reduced flexibility of loops at positions 82-91 and 207-215 from MM MD simulations.
